# Endogenous recruitment of frontal-sensory circuits during visual discrimination

**DOI:** 10.1101/2021.10.15.464378

**Authors:** Eluned Broom, Vivian Imbriotis, Frank Sengpiel, William M. Connelly, Adam Ranson

## Abstract

A long-range circuit linking anterior cingulate cortex (ACC) to primary visual cortex (V1) has been previously proposed to mediate visual selective attention in mice during visually guided behaviour. Here we used *in vivo* two-photon functional imaging to measure endogenous activity of ACC neurons projecting to layer 1 of V1 (ACC-V1_axons_) in mice either passively viewing stimuli or performing a go/no-go visually guided task. We observed that while ACC-V1_axons_ were recruited under these conditions, this was not linked to enhancement of neural or behavioural measures of sensory coding. Instead, ACC-V1_axon_ activity was observed to be associated with licking behaviour, modulated by reward, and biased towards task relevant sensory cortex.

## Introduction

Sensory processing is powerfully modulated by contextual and behavioural factors such as prior experience, anticipation, attention and movement (Ayaz et al., 2013; Keller et al., 2017, 2012; Leinweber et al., 2017; Morimoto et al., 2021; Niell and Stryker, 2010; Pakan et al., 2018; Poort et al., 2015; Ranson, 2017; Saleem et al., 2013, 2018; Speed et al., 2020). One mechanism of this modulation is thought to be long range glutamatergic cortico-cortical circuits (Gilbert and Li, 2013; Morimoto et al., 2021). Axons originating in higher cortical regions, such as frontal, retrosplenial and parietal cortex, and terminating preferentially in layer 1 of early sensory areas such as V1 are thought to influence sensory processing through both direct excitatory connections onto the tuft dendrites of V1 pyramidal neurons and indirectly through several classes of inhibitory neurons which in turn modulate excitatory neurons (Makino and Komiyama, 2015; Zhang et al., 2014).

A recent study (Zhang et al., 2014) identified one such circuit which connects the anterior circulate cortex to the primary visual cortex and demonstrated that it exerts an influence on sensory processing which shares similarities with selective visual attention described in primates (Armstrong et al., 2006; Moore and Armstrong, 2003; Reynolds and Heeger, 2009; Sundberg et al., 2009). Specifically, optogenetic activation of cingulate axons in V1 was shown to enhance the specificity of V1 neuron orientation tuning and to improve behaviourally measured visual discrimination. While this study demonstrated the anterior cingulate cortex → V1 projection could in principle function to enhance sensory processing, the relevance of the artificially induced patterns of cingulate activation to normal physiological function remains unclear. In particular, direct evidence of endogenous activity of V1 projecting ACC axons (ACC-V1_axons_) being linked to improved behavioural or neuronal stimulus discrimination is lacking. Subsequently other studies have argued axons originating from overlapping cingulate regions are selectively activated at a population level following errors during a freely moving 5-choice serial reaction time task (Norman et al., 2021a, 2021b), mediate sensory motor integration (Huda et al., 2020) and relay locomotion driven motor signals to V1 (Leinweber et al., 2017).

Here we aimed to disambiguate the attentional function of this circuit by reproducing the go/no-go head fixed behavioural paradigm employed in Zhang et al. 2014 but measuring at single axonal bouton resolution the endogenous recruitment of ACC-V1_axons_, and their relationship to behavioural and neural stimulus discrimination. In this context we find no evidence of an association between endogenous recruitment of this circuit and enhanced behavioural or neural measures of stimulus discrimination. Instead, we unexpectedly observe strong endogenous recruitment of the circuit in a subset of boutons by licking motor behaviour that is modulated by reward and biased towards the hemisphere currently processing task relevant sensory signals.

## Results

We first sought to assess previous claims of a role of long-range cortico-cortical projections from the cingulate (Cg) to the primary visual cortex (V1) in exerting topdown modulation of V1 activity which can enhance behavioural and neural visual discrimination accuracy (Zhang et al., 2014). While optogenetic activation of ACC-V1_axons_ during a go/nogo orientation discrimination task was previously shown to enhance discrimination accuracy, we aimed to evaluate if this circuit is also endogenously recruited in this way during visual processing.

### Cg → V1 axons are endogenously recruited during visual discrimination

ACC-V1_axons_ were labelled using the genetically encoded calcium indicator GCaMP6f and imaged using 2-photon microscopy in layer 1 of V1 (Figure 1A, B and S1A) in animals trained to perform a go/nogo stimulus orientation discrimination task at a high level of accuracy (discrimination as quantified by d’ > 1.5). We first tested whether the activity (ΔF/F) of ACC-V1_axons_ differed between within-trial periods (when the animal was actively engaged in discrimination) and inter-trial periods. We found that activity was significantly higher during discrimination vs. intertrial periods in 14% of ACC-V1_axons_ (Figure 1C and D; 84/597; paired sample t-test; see methods; summarized with task modulation index in Figure 1E), suggesting that the go/nogo behaviour results in recruitment of this circuit. Unexpected a further 21% of ACC-V1_axons_ (Figure 1C and D; 126/597) exhibited the opposite behaviour of higher levels of activity during intertrial periods suggesting suppression by task engagement.

**Figure 1.**
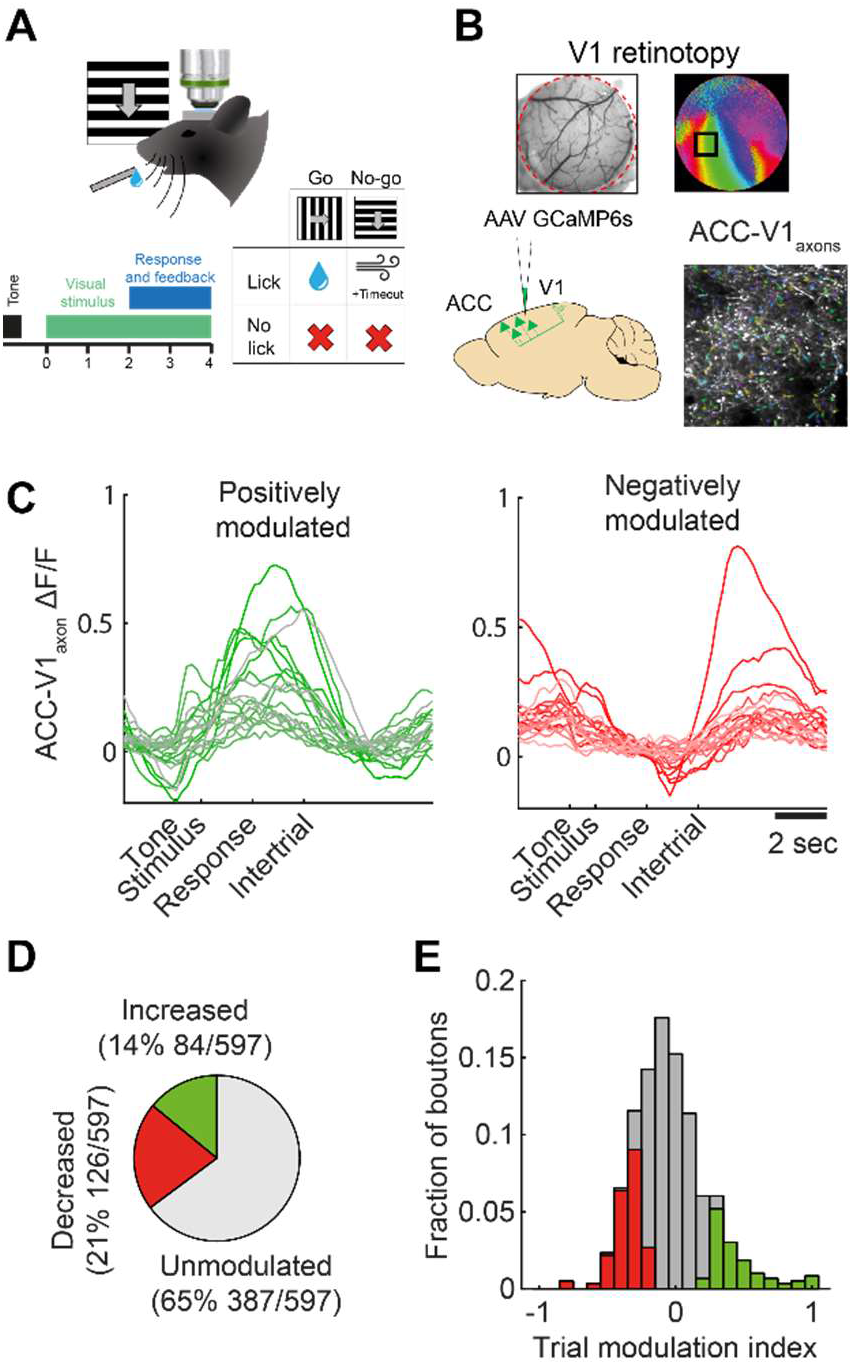
ACC-V1_axons_ are endogenously recruited during go/no-go visual discrimination behaviour. (A) Schematic of monocular go/no-go visual discrimination task. (B) Visualisation of the location of monocular primary visual cortex using intrinsic signal imaging and labelling of ACC-V1_axons_ by injection in cingulate cortex followed by visualisation using *in vivo* 2-photon microscopy in monocular V1 after 2-4 weeks. (C) Example boutons which were positively (green) or negatively (red) modulated in vs. out of trial. (D) In vs. out of trial modulation behaviour of all boutons. (E) Distribution of in vs. out of trial modulation indices of all boutons with statistically significant boutons indicated with colored bars.

### Endogenous ACC-V1_axons_ activity does not co-vary with behaviourally reported visual discrimination

Given previous findings of enhanced visual discrimination with artificial activation of ACC-V1_axons_ (Huda et al., 2020; Zhang et al., 2014), we next asked whether enhanced discrimination accuracy was also associated with endogenously elevated activity of ACC-V1_axons_. Within each behavioural session there was slow variation in accuracy over timescales of minutes (i.e. fluctuations in hit rate and false alarm rate and consequently d’; Figure 2A and B) suggesting varying levels of arousal, attention or task engagement. To test the association between ACC-V1_axon_ activation and this fluctuating discrimination accuracy, we fit a linear model to predict average bouton activity in each trial, using trial correctness and trial type (go or nogo) as predictors. A statistically significant association between ACC-V1_axons_ activity and trial correctness was observed in only a small fraction of boutons (5.7%, 34/597 ACC-V1_axons_, from 5 behavioural sessions from 5 mice; Figure 2C), and of these, while 47% were more active during correct trials, 53% were less activity during correct trials (Figure 2C inset). These results suggest that endogenous activation of ACC-V1_axons_ is unlikely to be playing a significant role in enhancing visual discrimination in this behavioural context.

**Figure 2.**
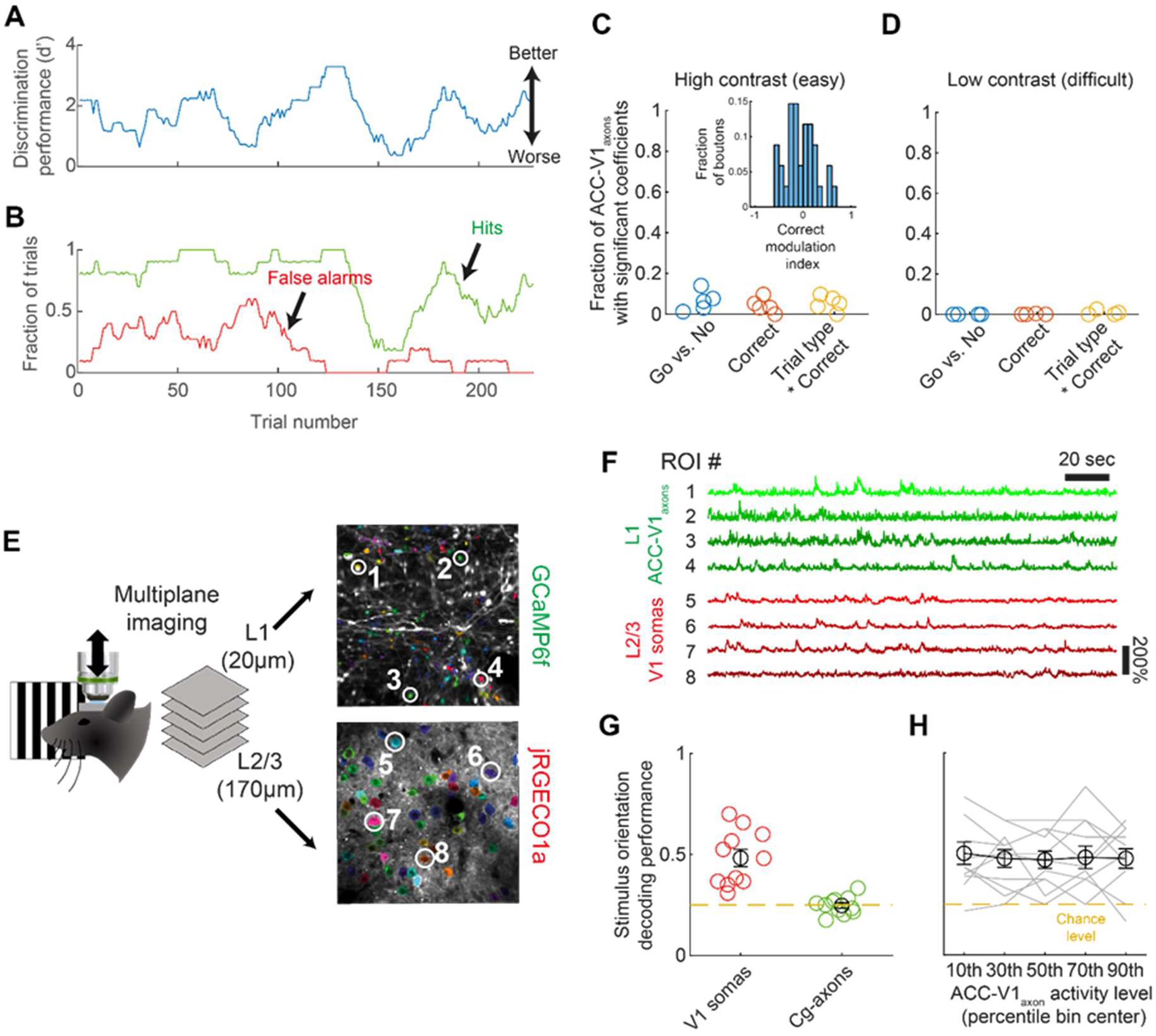
Behavioural and neural sensory discrimination accuracy are not enhanced by increased ACC-V1_axons_ activity. (A) Discrimination accuracy fluctuates markedly within each behvioural session as quantified by d’. (B) Discrimination accuracy fluctuation is driven both by variation in hit and false alarm rates. (C) Behavioural accuracy (correctness) is not associated with level of activity of ACC-V1_axons_ (i.e. ΔF/F) under high contrast (i.e. easier) stimulus conditions. The majority of the small fraction of boutons with an association between correctness and ΔF/F exhibit a negative association of higher accuracy with lower level of ACC-V1_axon_ activity. (D) As in (C) but at low stimulus contrast and with comparable findings. (E) Schematic of multiplane imaging to concurrently record cingulate axons in layer 1 (labelled green with axon-GCaMP6f) and V1 neurons in layer 2/3 (labelled red with jRGECO1a). (F) Example traces of concurrently recorded boutons (green) and somas (red). (G) Cross validated accuracy of a SVM classifier decoding stimulus orientation from the activity of V1 somas or ACC-V1_axons_. (H) Classifier accuracy decoding visual stimulus orientation at different levels of ACC-V1_axon_ population activity.

In the trials analysed above the stimulus contrast was maximum and discrimination accuracy was high (mean d’ = 1.67). We hypothesised that this could cause a ceiling effect whereby improvements in stimulus discrimination due to top-down modulations in the efficacy of V1 encoding are limited, or that the circuit may only be recruited when task demands are high (Bahrami et al., 2007; Norman et al., 2021b). We tested this possibility in a different group of animals by increasing task difficulty by altering stimulus contrast in a subset of trials. As expected discrimination accuracy decreased at lower contrasts (Figure S1B; high contrast [50%] d’ = 1.89 ± 0.12; low contrast [10%] d’ = 1.11 ± 0.05; n = 4 mice; p = 0.001; paired sample t-test). We again tested whether cingulate-axon activity was elevated on correct vs incorrect trials when the task was more difficult. This analysis also showed that almost no cingulate-axons (<0.1%) showed activity which differed significantly between correct and incorrect trials. We assessed whether behavioural errors might instead drive recruitment of ACC-V1_axons_,(Norman et al., 2021a) however ACC-V1_axon_ activity was also not found to differ significantly as a function of previous trial correctness (see Figure S4; and Supplemental results). Together these results indicate the lack of association between endogenous cingulate axon activity and discrimination accuracy is unlikely to be due to task difficulty in the context of this task.

### Endogenous ACC-V1_axon_ activity does not co-vary with neurally measured stimulus discrimination

Optogenetic activation of ACC-V1_axons_ has also been shown to enhance orientation tuning in V1 during passive viewing (Zhang et al., 2014). These findings prompted us to ask if V1 encoding might be improved by endogenous activity of ACC-V1_axons_ even in the absence of behaviour enhancement. In support of this possibility, a disassociation has been reported between visual stimulus encoding fidelity in V1 in mice and behavioural readouts of visual discrimination, whereby neural encoding precision of sensory stimuli significantly exceeds that measured behaviourally (Stringer et al., 2021). We used multiplane imaging and red and green calcium indicators to concurrently measure activity of ACC-V1_axons_ in layer 1 (labelled green with GCaMP6s) and V1 neuron somas (labelled red with jRGECO1a) in layer 2/3 during stimulation with drifting gratings (Figure 2E and F). To assess visual stimulus encoding in V1 we trained a support vector machine classifier to decode visual stimulus orientation from the activity of populations of V1 neurons, with the accuracy level of the classifier determined using leave-one-out cross validation. As expected, stimulus orientation could be decoded at above chance level from activity of V1 neurons (n = 11 fields of view, from 3 mice, with an average of 103±14 cells per field of view; mean accuracy 48.24±4.31% correct; chance level = 25%; t-test against chance level: p = 0.0002; Figure 3C). Note that decoding accuracy was limited due to the number of V1 neurons that could be recorded at the relatively high level of zoom required to concurrently record axonal boutons (see Figure S2A for quantification). In contrast, stimulus orientation could not be reliably decoded from the ACC-V1_axon_ populations recorded (n = 11 fields of view, from 3 mice, with an average of 192±27 boutons per field of view; mean accuracy 24.72±1.37; t-test against chance level; p = 0.83; Figure 2G) suggesting orientation information is not relayed through this feedback circuit.

**Figure 3.**
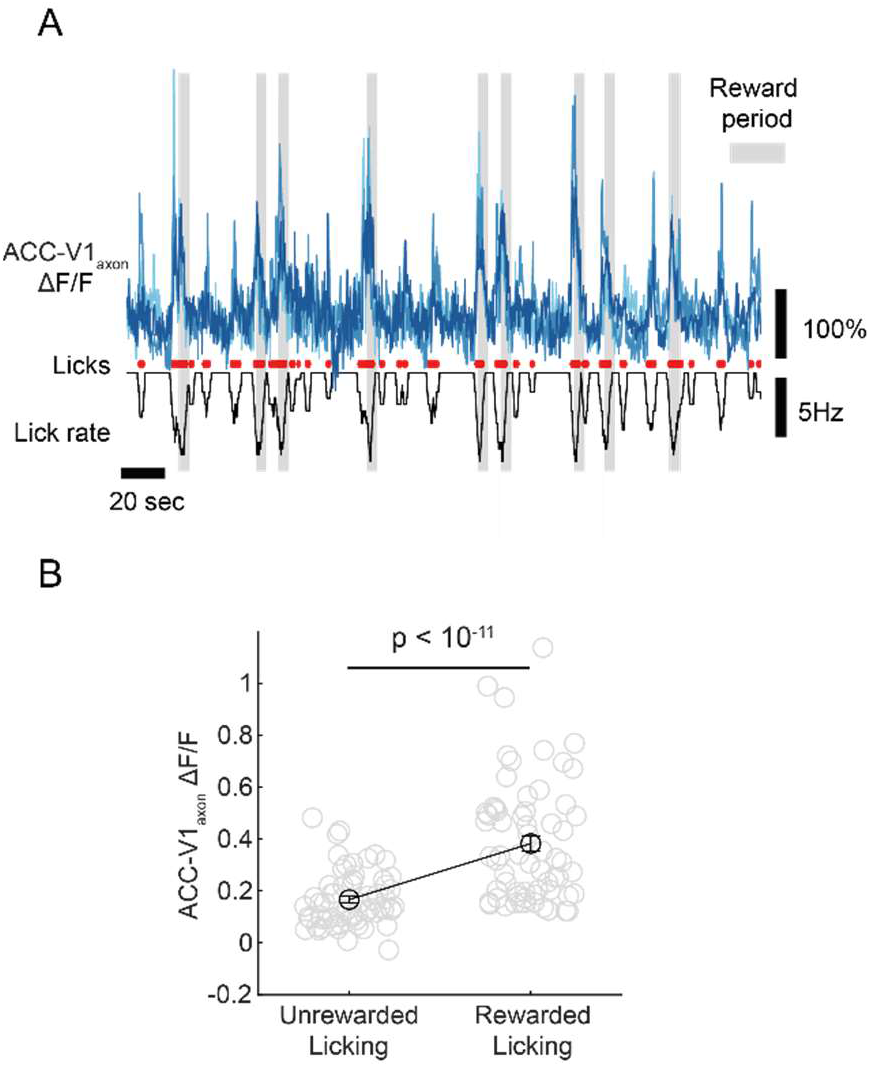
Licking associated activity of ACC-V1_axons_ is modulated by reward. (A) Example traces of lick correlated boutons (shades of blue), individual lick events (red) and lick rate (black; note more negative is higher lick rate). Gray areas show periods when licking is rewarded. (B) In lick correlated boutons, rewarded licking results in a higher level of activity (ΔF/F) than unrewarded licking.

We next tested whether classifier orientation decoding accuracy increased with increased endogenous population ACC-V1_axons_ activity as might be anticipated from previous findings of artificially activating this feedback pathway. Mean population activity of ACC-V1_axons_ varied significantly during the course of each experiment (64±0.04% difference between 20th and 80th percentile), allowing us to test the accuracy of the decoder at predicting stimulus orientation from the activity of V1 neurons at different levels of ACC-V1_axon_ population activity. The decoder accuracy was assessed for each trial using a model trained on all other trials. The trials within each recording session were then divided into 5 sets based on the level of activity of the ACC-V1_axon_ population during each trial set borders were defined as the 20th, 40th, 60th, 80th percentiles of ACC-V1_axon_ population activity over the entire recording session. An average of the classifier accuracy for the trials within each set was then calculated. We found no relationship between decoder accuracy and ACC-V1_axon_ activity level (Figure 2F; 1-way ANOVA; n = 11 experiments from 4 mice; p = 0.99).

We considered whether different results might be obtained by training and testing the classifier using only trials from within each ACC-V1_axon_ activity level set (instead of the entire session) but results were comparable (Figure S2C). Consistent with these findings, a comparison of the orientation tuning curves of individual V1 neurons measured during high or low levels of ACC-V1_axon_ activity (less than vs. greater than 50th activity percentile) showed that orientation preference and selectivity did not differ systematically in the population overall (Figure S2C and D), although in individual cells apparent alterations could be observed (Figure S2E and F). Overall these findings are inconsistent with the idea that endogenous fluctuations in ACC-V1_axon_ activity alter orientation encoding in V1.

### ACC-V1_axon_ activity is associated with rewarded licking

A subset (17%) of ACC-V1_axons_ showed statistically significant licking correlated activity which explained some of the recruitment of these axons during the discrimination phase of the task (tested with permutation test with 1000 shuffled lick rate traces; p < 0.05; Figure 3A). As licking often co-occurred with reward we sought to disambiguate these two factors. Licking was often not rewarded (i.e. during inter-trial periods and in no-go trials) allowing a comparison of ACC-V1_axon_ recruitment between rewarded vs. non-rewarded licking. We compared the fluorescence (ΔF/F) of highly lick correlated boutons (R>0.2) during rewarded licking (within go trial reward period) versus unrewarded licking (during inter-trial periods) and found that almost all boutons were more active during rewarded vs. unrewarded licking, suggesting licking associated ACC-V1_axon_ activity encodes a combination of both licking and reward (unrewarded vs. rewarded ΔF/F = 0.17 ± 0.01 vs. 0.38 ± 0.03; p < 10^−11^; paired sample t-test; Figure 3B).

Rewarded licking patterns tended to be more long lasting and rhythmic (typically >6Hz) than non-rewarded licking which could result in a difference in neural activity unrelated to reward per se. To control for this, we searched for periods of unrewarded rhythmic licking at >6Hz. Unrewarded licking bouts were typically shorter than bouts of rewarded licking and so we limited analysis to ACC-V1_axon_ activity during the first 3 licks in a bout of 3 or more licks. Consistent with the previous analysis, we found that frequency and count matched licking bouts were associated with greater ACC-V1_axon_ activity when rewarded than unrewarded (unrewarded vs. rewarded ΔF/F = 0.19 ± 0.01 vs. 0.37 ± 0.029; p < 10^−8^; paired sample t-test; Figure S3). This result indicates differences in rewarded versus unrewarded licking are unlikely to be explained by licking pattern. These data suggest that ACC-V1_axon_ activity encodes in part a form of reward signal.

### ACC-V1_axon_ licking/reward signals are biased towards task relevant sensory cortex

We next asked if the licking/reward signals observed are target to the areas of the visual cortex processing the sensory signals that are guiding discrimination behaviour or alternatively are also relayed non-specifically to areas not involved in the discrimination. We trained animals on a variant of the go/no go task (Figure 4A) in which the stimulus was presented randomly to either the contralateral or ipsilateral monocular visual field such that on each trial the stimulus was being primarily processed in V1 in either the imaged hemisphere (hemisphere contralateral to the stimulus) or V1 in the non-imaged hemisphere (hemisphere ipsilateral to the stimulus). Animals learned the bilateral version of the task to a high level of accuracy and tended to have a similar level of accuracy when processing visual stimuli via the left and right hemispheres (Figure 4B and C). Fluctuations in task accuracy was most often correlated between the left and right monocular visual field discrimination trials (suggesting a global cause of fluctuation of accuracy; Figure 4C), however sometimes fluctuations were side dependent suggesting possible hemisphere specific effects (Figure S3B). As observed in the monocular version of the task, a subset of ACC-V1_axons_ showed activity correlated with licking (Figure 4D), and of these almost all showed greater activity during rewarded vs. non-rewarded licking on both contralateral and ipsilateral stimulus trials (Figure 4E). To test the specificity of the hemispheric targeting of these licking/reward signals we examined if, on a trial-by-trial basis, rewarded licking signals were preferentially targeted to V1 in the hemisphere where task related sensory signals were primarily arriving (i.e. the hemisphere contralateral to the monocular stimulus) or alternatively to V1 in both hemispheres. To quantify this, we calculated a ‘reward targeting index’ (RTI) whereby values of −1 and 1 respectively indicate reward signals are exclusively targeted to V1 ipsilateral or contralateral to the stimulus, while a value 0 indicates equal targeting to the two hemispheres. The median RTI was 0.09 (Figure 4F; 64% of boutons had an RTI > 0; one sample t-test for difference from 0; p = 0.002) indicating a modest bias in reward signal targeting to V1 in the hemisphere processing the task related stimulus. This was reflected in a mean rewarded ΔF/F of 0.63 ± 0.04 vs. 0.47 ± 0.04 in the contralateral vs. ipsilateral hemisphere (p = 0.004; paired t-test).

**Figure 4.**
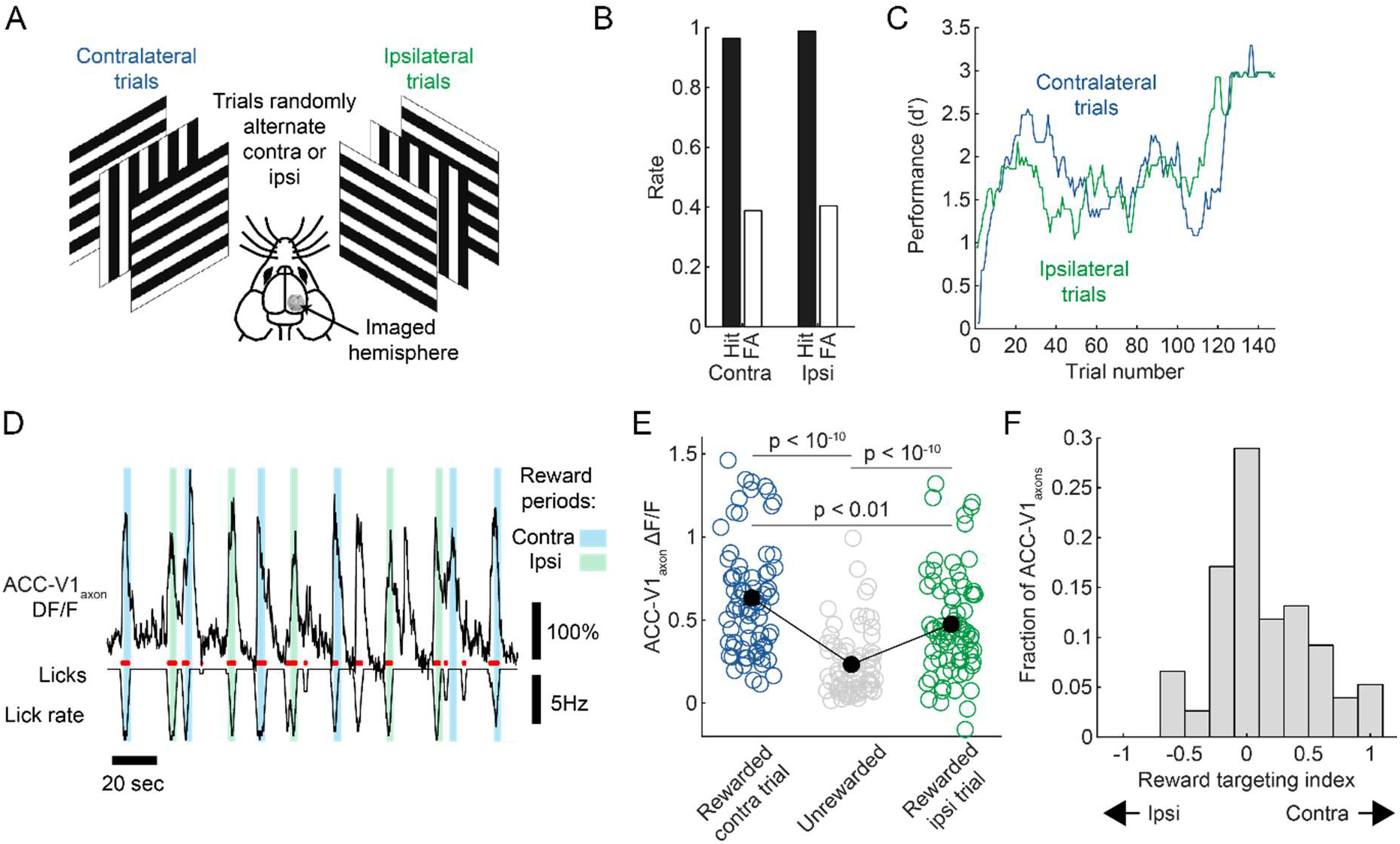
Licking associated activity in ACC-V1_axons_ is modulated by reward and hemispherically biased depending upon current task conditions. (A) Schematic of bilateral version of the visual discrimination task. (B) accuracy is typically similar on ipsilateral and contralateral trials. (C) During the course of one experiment contralateral and ipsilateral accuracy often vary together. (D) Example trace of lick correlated bouton in bilateral task, individual lick events (red) and lick rate (black; note more negative is higher lick rate). Light blue and light green shaded areas show contralateral and ipsilateral trial reward periods respectively. (E) In lick correlated boutons rewarded licking results in a higher level of activity (ΔF/F) than unrewarded licking, and this effect is greatest on contralateral trials (i.e. when the stimulus is being represented in monocular V1 in the same hemisphere as the ACC-V1_axons_ are originating from). (F) Distribution of ‘reward targeting index’ values.

## Discussion

Our experiments reveal that while ACC-V1_axons_ projecting to V1 are endogenously recruited during visual discrimination behaviour, this recruitment is not associated with enhanced behavioural discrimination accuracy. Instead we unexpectedly find a significant fraction of ACC-V1_axons_ exhibit activity which is strongly correlated with licking. In these boutons licking activity was modulated by whether it was rewarded and additionally moderately biased towards the hemisphere receiving task associated sensory data.

Our findings are at odds with some previous studies in which exogenous optogenetic excitation (Zhang et al., 2014) or suppression (Huda et al., 2020) of the same ACC → V1 projection has been found to respectively enhance orientation discrimination behaviour or impair visual detection driven orienting behaviour. Similarly, optogenetic excitation of this circuit has previously been reported to enhance orientation tuning in V1 during passive viewing (Zhang et al., 2014), while in our experiments endogenously heightened activity in the ACC-V1_axons_ population was not associated with a higher accuracy of orientation decoding from neural activity. In this respect our findings are more consistent with another report (Norman & Riceberg et al. 2021) in which fiber photometry was used to measure sum neural activity of cingulate neurons projecting to V1 in freely moving mice performing a 5-choice serial reaction time task. This study reported that while cingulate activity was found to be enhanced following errors, this did not result in improved behavioural accuracy. Together these findings suggest the possibility that the powerful and synchronous activation/inactivation of this circuit obtained using optogenetic approaches, and the consequent effects on behaviour and sensory coding, may be quite different from the range of activity this circuit exhibits endogenously.

We examined our dataset for evidence of previously reported post-error recruitment of V1 projecting cingulate neurons (Norman et al., 2021a), but found that even in the more demanding versions of our task there was no evidence for such a pattern of activity (posterror recruit of the circuit has been reported to be limited to conditions of higher task difficulty; Norman, Bateh, et al. 2021). One possibility is that our task is not sufficiently cognitively demanding, although we observed a clear reduction in accuracy when reducing stimulus contrast, suggesting we are not at some ceiling accuracy level. There are also likely to be important differences due to details of the specific task such as whether the animal is freely moving. It would be of interest to develop head-fixed task variants which reproduce the post-error circuit recruitment described by Norman & Riceberg et al. but with greater control of visual stimulus and motor behaviour than that possible in freely moving animals.

Other studies have described motor related activity in ACC-V1_axons_ (Huda et al., 2020; Leinweber et al., 2017), although not to the best of our knowledge licking/reward driven activity. Interestingly, Leinweber et al. studied V1 projecting axons from a likely overlapping region they identify as A24b and argued for a seemingly completely distinct role of the projection in relaying predictions to V1 of visual flow based on motor output. While we have observed clear motor related activity driven by licking, it is difficult to see what visually might be predicted to follow as a consequence of licking behaviour (beyond the offset of the stimulus at the end of the trial), and how this relates to the apparent reward contingency of this activity.

A number of previous studies have identified reward (Pakan et al., 2018; Poort et al., 2015; Schuler and Bear, 2009) related activity in V1 neurons, however the route through which these signals impinge upon V1 has remained unclear. Here we report ACC-V1_axons_ as a novel circuit through which reward and licking signals enter V1 in a way which is biased, at least at a gross scale, towards currently engaged regions of sensory cortex. The role that such reward and licking signals play in visually guided behaviour remains unclear, but we speculate that they may be involved in driving and maintaining previously described plasticity processes that result in alterations of the representations of previously rewarded sensory stimuli (Poort et al., 2015). Further experiments in which such reward signal carrying ACC-V1_axons_ are selectively inactivated will be required to test this possibility.

## Methods

### Animals

All experimental procedures were carried out in accordance with institutional animal welfare guidelines, and licensed by the UK Home Office. Experiments were carried out on adult mice (aged >P90). Mice were housed under normal light conditions (14h light, 10h dark) and recordings were made during the light period.

### Animal surgical preparation and virus injection

Aseptic surgical procedures were conducted based on previously described protocols (Goldey et al., 2014; Ranson, 2017). Approximately one hour prior to cranial window surgery and virus injection, animals were administered with the antibiotic Baytril (5mg/kg, s.c.) and the anti-inflammatory drugs Carprofen (5mg/kg, s.c.) and Dexamethasone (0.15mg/Kg, i.m.). Anaesthesia was induced and maintained using Isoflurane at concentrations of 4%, and 1.5-2% respectively. After animals were stereotaxically secured, the scalp and periosteum were removed from the dorsal surface of the skull, and a custom head plate was attached to the cranium using dental cement (Super Bond C&B), with an aperture approximately centred over right V1. A 3mm circular craniotomy was next performed, centred on the stereotaxically identified monocular area of V1.

For injections into V1 intrinsic signal imaging was used (after skull exposure but before craniotomy) to localise monocular V1 (see methods below for description of intrinsic signal imaging) after which injections were targeted using a functional map of retinotopy overlaid on surface vasculature (depth 250μm). After craniotomy an AAV was injected to drive expression of jRGECO1a (AAV1.Syn.NES-jRGECO1a.WPRE.SV40; titre after dilution 5×10^12^ GC/ml; volume 100nl). For injections into ACC a small craniotomy was first made over the region (centred at 0.2 - 0.3 mm anterior and 0.3 lateral of bregma) either using a dental drill or by thinning the overlying bone and then piercing a small hole using a hypodermic needle. After craniotomy an AAV was injected to drive expression of GCaMP6s (AAV1.Syn.GCaMP6s.WPRE.SV40; titre after dilution 2×10^11^ GC/ml; volume 100nl; used in behavioural experiments) or axon-GCaMP6s (rAAV2/1-hSynapsin1-axon-GCaMP6f; titre after dilution 5×10^11^ GC/ml; volume 100nl; used in passive visual stimulation experiments (Broussard et al., 2018)). All injections were made using a Nanoject II system at a rate of 10nl/minute using pulled and bevelled oil filled glass micropipettes with a tip outer diameter of approximately 30μm. After injections the craniotomy over V1 was closed with a glass insert constructed from 3 layers of circular no 1 thickness glass (1×5 mm, 2×3 mm diameter) bonded together with UV cured optical adhesive (Norland Products; catalogue no. 7106), and the craniotomy over ACC was closed with Vetbond. After surgery animals were allowed at least 2 weeks to recover after which they were either habituated to head fixation during passive visual stimulation or during visual discrimination training.

### In vivo imaging

*In vivo* 2-photon imaging was performed using a resonant scanning microscope (Thorlabs, B-Scope) with a 16x 0.8NA objective with 3mm working distance (Nikon). Genetically encoded calcium indicators were excited at 920-980nm using a Ti:sapphire laser (Coherent, Chameleon) with a maximum laser power at sample of 50mW. Data was acquired at a framerate of approximately 60Hz and averaged, resulting in a framerate of approximately 10Hz. Imaging, behavioral and visual stimulation timing data were acquired using Scanimage 4.1 and custom written code (Matlab) and a DAQ card (NI PCIe-6323, National Instruments). In *vivo* intrinsic signal imaging was performed using previously described methods(27) using either a custom built system based around a MAKO G-125B camera (AVT) or a commercially available system (Imager 3001, Optical Imaging Inc.).

### Passive visual stimuli

For recordings of passive visual responses (i.e. not during behaviour) mice were stimulated with a circular 20×20 deg drifting horizontal square wave gratings with temporal frequency of 2 Hz and spatial frequency of 0.05 cycles per degree, and at one of 8 orientations. The stimuli were displayed at one of 32 positions arranged in a grid of 8 horizontal positions (spanning 80 deg of visual space) and 4 vertical positions (spanning 30 deg of visual space). Each stimulus appeared and drifted for 1 second after which the next stimulus was displayed. Visual stimuli were generated in Matlab using the psychophysics toolbox (Brainard, 1997) and displayed on calibrated LCD screens (Iiyama, BT481).

### Behaviour

Animals were trained in a go/no-go task similar to that previously described (Andermann et al., 2010; Zhang et al., 2014), and implemented using custom written code in Matlab using the psychophysics toolbox (Brainard, 1997). In the task animals had to lick in response to a horizontally oriented drifting grating for water reward (go condition), or suppress licking in response to a vertically oriented drifting grating (no-go condition). Grating stimuli occupied approximately 80 deg of visual field and were presented in the monocular visual field. Each correct trial was rewarded with 5μl of a solution made up of 500ml water, 50g sucrose and 1.7g Koolaid. Licking was detected using a custom made capacitive lick sensor. Rewards were delivered using a reward valve (Neptune Research 161T011) controlled using a custom made circuit triggered with a digital signal of calibrated duration from a data acquisition device (Labjack, U3) interfaced with using Matlab. Incorrect trials were punished with an air puff, white noise auditory stimulus and a ten second time-out. Several days prior to commencing training, mice were placed on water restriction and behavioural training commenced after they reached approximately 80% of their initial weight. During the initial stage of training no visual stimuli were presented to the mice and animals could obtain a reward by licking during windows of up to 60 seconds. If animals licked during this period they were rewarded and this was followed by a variable duration period (2 - 10 secs) during which licking was not rewarded. During the variable delay period (termed the quiescent period) animals had to suppress licking or the next trial did not start. This quiescent period was maintained during the entire training and testing procedure. Once mice were consistently licking the spout they progressed to the next stage during which trials were initiated with an auditory tone, followed by 4 seconds of go stimulus presentation. The mouse was rewarded for licking during the 2 to 4 second period after go-stimulus onset - i.e. responses during the initial 2 second period after stimulus onset were disregarded. Disregarding licks immediately after stimulus onset was important as often animals exhibited impulsive licking at stimulus onset which was unrelated to stimulus type. During this training period, in a gradually decreasing fraction of trials (starting at 90% free trials), free rewards were administered during the go stimulus period, even if the animal didn’t respond. This fraction of free reward trials was automatically decreased in steps of 10% if animals responded independently and correctly in blocks of 20-30 trials. Once animals were responding correctly to 70% of go stimuli the no-go stimulus was introduced and free reward trials were excluded. Once animals reached a go/no-go discrimination accuracy of d’ > 1.5 they were classified as fully trained and at his point experimental imaging data was typically collected. The discriminination index d’ was calculated in Matlab using norminv(hit rate)-norminv(false alarm rate). In the ‘bilateral’ version of the task, in each trial stimuli were presented in either the left or right monocular visual field, but the training criteria procedures were as above. In the bilateral task, training progress was measured, and training stage advanced, using overall accuracy (i.e. pooled over left and right trial types) rather than in a side specific manner.

### Calcium imaging data pre-processing

Calcium imaging data was registered and segregated into neuronal regions of interest (i.e. Cg-boutons or V1 somas) using Suite2P (Pachitariu et al., 2016). The time series of each ROI was then converted from a raw fluorescence value to dF/F with the denominator F value trace constructed by calculating the 5th percentile of the smoothed F value within a 20 second window centred on each sample in the F trace. Boutons with correlation coefficients of > 0.7 were considered to be from the same axon and combined using a weighted average, with weighting determined by the number of pixels in the ROI of the bouton.

### Analysis of activity of ACC-V1_axons_ during behaviour

Whether boutons were overall positively or negatively modulated by the task (Figure 1D-F) was calculated by comparing within task periods (average ΔF/F value over the period from trial onset tone to the stimulus offset) to between task periods (average ΔF/F value of the final 2 seconds of the properly executed quiescent period immediately before trial onset; thus ensuring a period with no licking behaviour). The task modulation index (Figure 1F) was calculated from these same periods with the formula (within task activity - between task activity) / (within task activity - between task activity) with an index of 1 indicating boutons are only active inside of the task and an index of −1 indicating boutons are only active during the intertrial quiescent period.

To assess the association between the activity of individual ACC-V1_axons_ and task accuracy (Figure 2C and D) we constructed a linear model to predict individual bouton activity in each trial, using trial correctness, trial type (go or nogo) and an interaction term as predictors. Bouton activity was calculated in each trial as the average ΔF/F value over the period from trial onset tone to the stimulus offset. The linear model was implemented using the fitglm function in Matlab with correction for multiple hypothesis testing (i.e. testing multiple axons) implemented using the mafdr false discovery rate function. The correct modulation index (Figure 2C inset) was calculated as (correct task bouton activity - incorrect task bouton activity) / (correct task bouton activity - incorrect task bouton activity).

To measure the correlation between licking and ACC-V1_axon_ activity the lick raster was first converted to a lick rate by summing the licks in a 1 second window which was slid over the binary lick event trace. The correlation coefficient between lick rate and the ΔF/F of each ACC-V1_axon_ roi was then calculated. Because of the nonindependence of neighbouring samples in the two traces we used a permutation test to test significance by building a null distribution to which the observed correlation coefficients could be compared. To do this we randomly circularly shifted the lick rate trace 1000 times, with a minimum shift of the equivalent of ±10 seconds, and in each instance measured the correlation coefficient between the shifted lick rate trace and ΔF/F of each ACC-V1_axon_ roi. ACC-V1_axons_ were deemed to be significantly correlated if their observed lick rate correlation coefficient exceeded 95% of those observed in the null distribution. The comparison of rewarded vs. unrewarded licking (Figure 4B and 4G) was made by averaging ΔF/F during rewarded licking within trial and unrewarded licking outside of trial. In analysis where we sought to control for licking frequency, bouts of licking were identified using the raw lick events, which were isolated in time (i.e. preceded by at least 2 seconds of non-licking) and only the neural activity during the first 3 licks in a bout were analysed to avoid confounds associated with rewarded licking bouts differing in length from unrewarded licks.

In order to analyse the hemispheric targeting of lick/reward signals a targeting index was derived from the mean ΔF/F during rewarded licking in contralateral trials (C_reward_) and ipsilateral trials (I_reward_) using the formula: (C_reward_ - I_reward_) / (C_reward_ - I_reward_).

### Analysis of activity of ACC-V1_axons_ during passive visual stimulation

SVM classifier analysis (Figure 3) was performed in Matlab by using the fitcecoc function to fit a multi-class support vector machine model using a one-versus-one coding design resulting in n(n - 1)/2 SVMs (i.e. 6 SVMs to decode which of 4 stimulus orientations was being displayed), using the default linear kernel and box constraint parameter of 1 (resulting in a relatively ‘soft’ margin). The average ΔF/F of each ROI (either V1 soma or ACC-V1_axons_) was taken over the 1 second after stimulus onset in each trial and each ROI constituted a predictor of the class labels (i.e. the stimulus orientations) in the model. Overall classifier accuracy (Figure 3C) was assessed using leave-one-out cross validation (i.e. multiple models were fit, each with one of the trials left out which was subsequently used for testing of the fitted model from which it was excluded during training), with the accuracy reported being the mean accuracy of all models. The level of activity of the ACC-V1_axon_ population was taken as a mean of all ACC-V1_axon_ rois detected. Orientation tuning data (Figure S2F) were fitted using the Matlab function lsqcurvefit and modelled as a sum of two Gaussians which were constrained such that one peaked at the preferred stimulus, and the peaks were 180 degrees apart (Carandini and Ferster, 2000).

### Other statistical Methods

Statistical analysis was carried out in MATLAB 2019b using the Statistics toolbox, and group average values are presented throughout as mean ± standard error of the mean. The statistical significance of comparisons between groups was determined using a two-sided t-test or ANOVA unless otherwise noted, and p values < 0.05 were considered significant. Similarity of variance and normal distribution were checked with the vartestn Matlab function. Correction of p values for multiple comparisons were calculated using the Matlab function multcompare using the Tukey–Kramer method. Precise group sizes were not decided in advance but approximate group sizes were based on typical sizes used in this field in similar experiments.

## Acknowledgements

This work was supported by a BBSRC funded PhD fellowship (SWBio DTP) to F.S. and A.R., a Sêr Cymru Fellowship (80762-CU-080) to A.R, a Wellcome Trust ISSF Seedcorn Award (105613/Z/14/Z) to A.R. and grant PDI 2019-109285GA-I00 from the Spanish Secretary of Research, Development and Innovation (MINECO) to AR.

## Supplemental information

### Supplemental Results

#### Post error recruitment of ACC-V1_axons_ does not occur in this task

One recent study reported V1 projecting cingulate neurons are selectively activated at a population level following errors during a freely moving 5-choice serial reaction time task (Norman et al., 2021a). We thus tested for this phenomenon in our dataset. We pooled all experiment types examined in the study (32 behavioural sessions in total) and compared cingulate axon population activity (average of all boutons imaged in each experiment) following correct or incorrect trials, both in the intertrial period and during the within trial stimulation period, but found no differences in population ΔF/F as a function of previous trial correctness (Figure S4A; two-way ANOVA; effect of previous trial p = 0.34). In a follow-up to the above study, a comparison of cingulate in 5-choice vs. 2-choice serial reaction time tasks revealed that cingulate circuit recruitment following errors depends upon task difficulty (Norman, Bateh, et al. 2021) with recruitment only occurring when task difficulty was higher (i.e. 5-choice version of task). Two variations of the task we used in our study are arguably higher difficulty, the variant where stimulus contrast is reduced, and the binocular variant where the animal does not know in advance where the stimulus will appear. We tested for prior trial correctness effects in these subsets of experiments but also observed no differences (low contrast: Figure S4B; two-way ANOVA; effect of previous trial p = 0.65; low contrast and binocular: Figure S4C; two-way ANOVA; effect of previous trial p = 0.71).

**Supplemental Figure 1.**
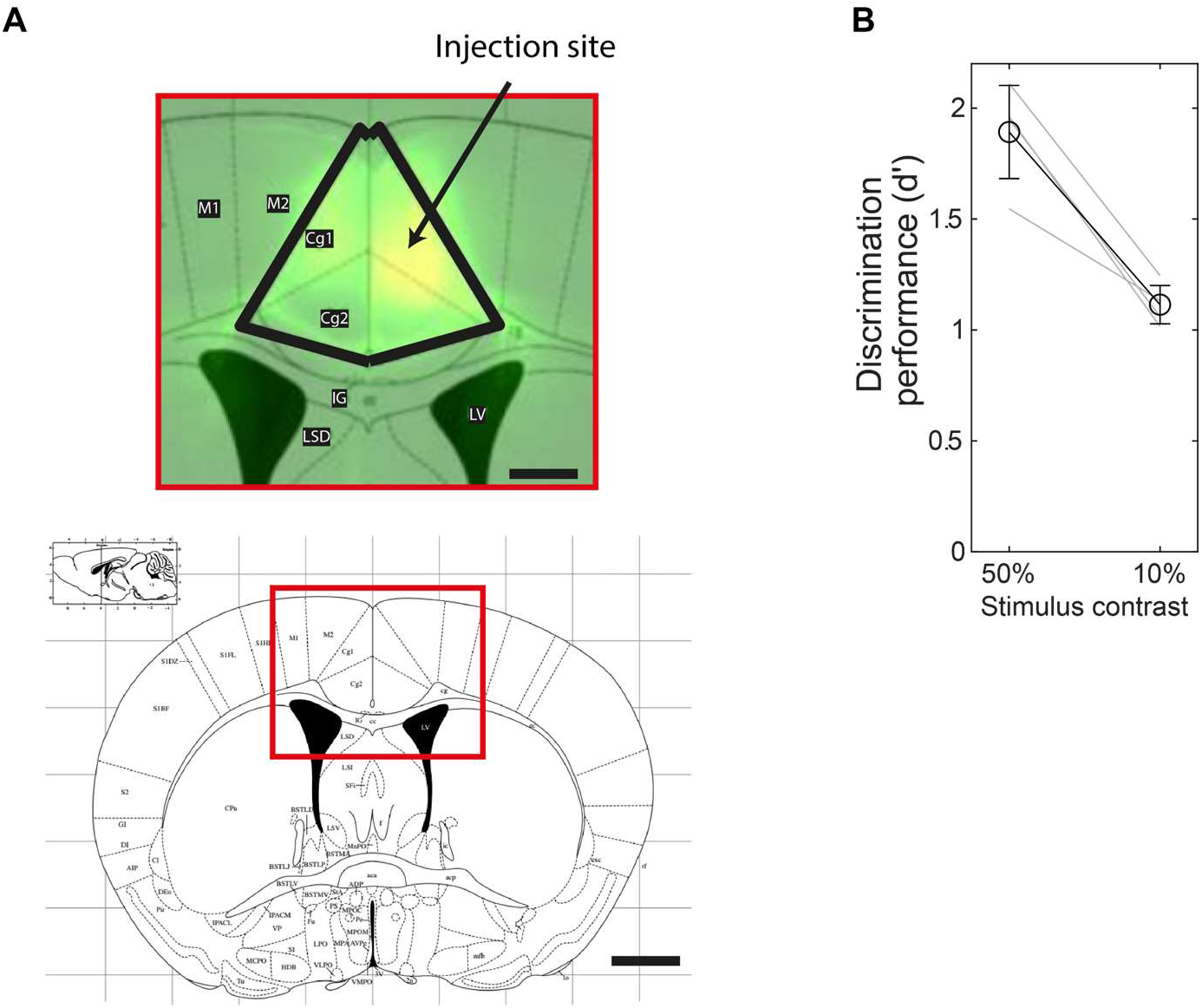
(A) Histology from ACC injection site (scale bars: upper panel = 0.5mm, lower panel = 1mm). Note, injection was unilateral, but ACC axons can be seen to project to contralateral ACC. (B) Discrimination accuracy reduces at lower compared to higher stimulus contrasts.

**Supplemental Figure 2.**
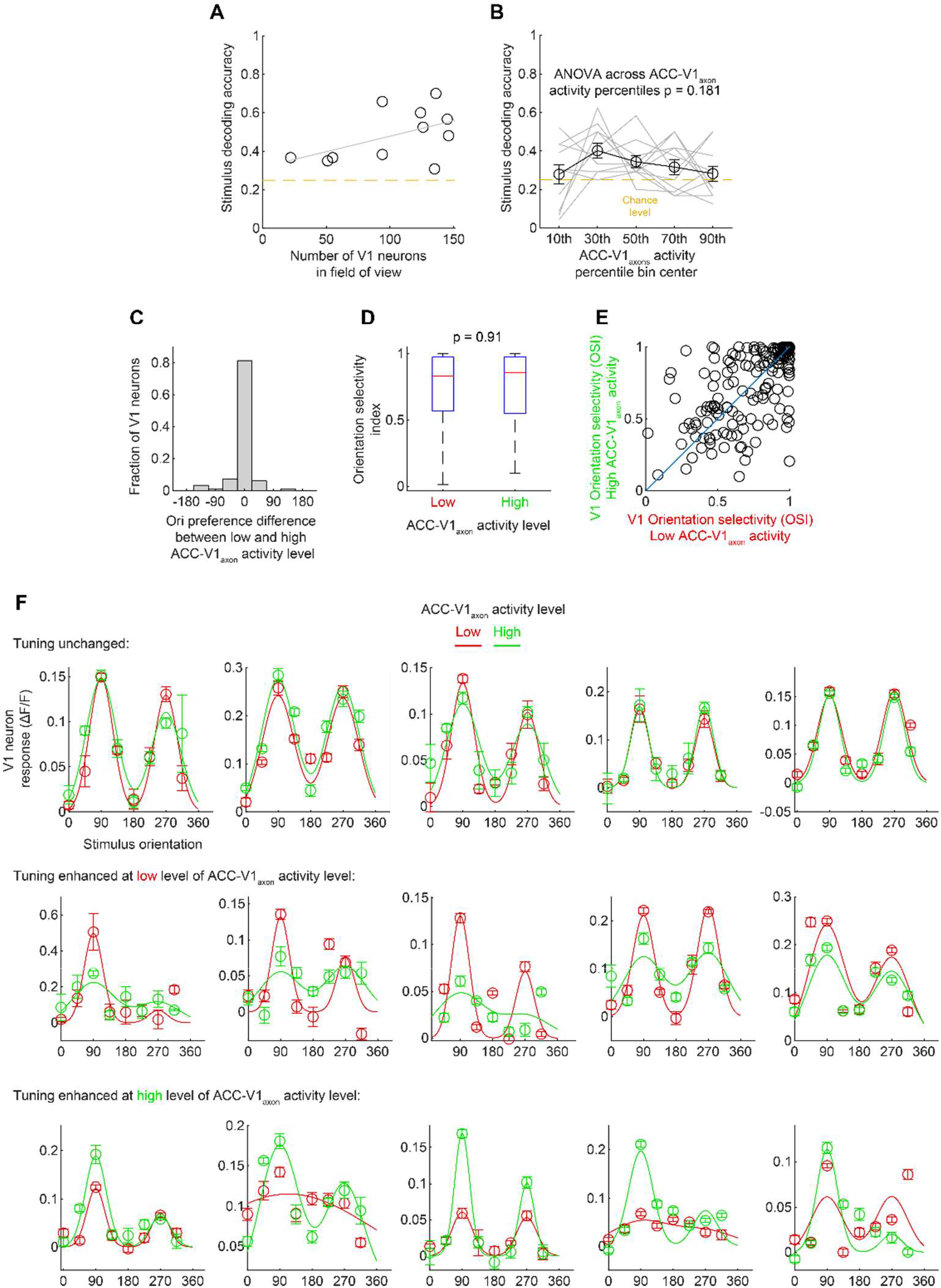
(A) SVM decoder accuracy increases with numbers of neurons detected in the field of view. (B) Results are similar to Figure 3D when the SVM is trained and tested using trials with the same ACC-V1_axon_ activity level. Overall orientation tuning preference (C) and orientation selectivity (D and E) does not on average differ significantly between trials with low compared to high levels of ACC-V1_axon_ activity. (F) Individual cells do exhibit shifts in orientation tuning selectivity as a function of increased ACC-V1_axon_ activity, but these shifts are not systematically towards enhancement or degradation of tuning. Comparison of OSI in (D) was made using a Kruskal–Wallis test (n = 192 cells, from 11 experiments from 3 mice). Analysis of orientation tuning curves was limited to cells which were classified as visually responsive (one-way ANOVA over all stimulus conditions), and for which the R^2^ of orientation tuning curve fits (at both low and high levels of ACC-V1_axon_ activity) was > 0.3.

**Supplemental Figure 3.**
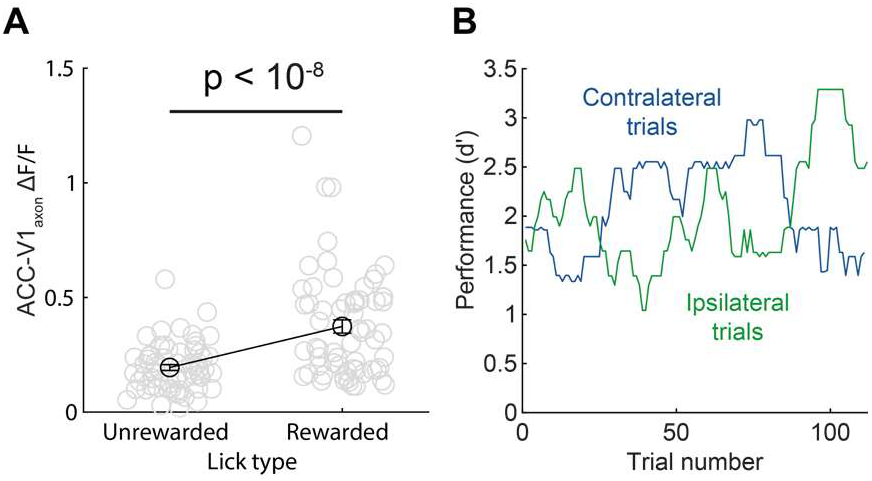
(A) Rewarded licks are associated with greater in ACC-V1_axon_ activity than unrewarded licks after licking pattern is controlled for. (B) In some experiments accuracy on ipsilateral vs. contralateral trials varies independently.

**Supplemental Figure 4.**
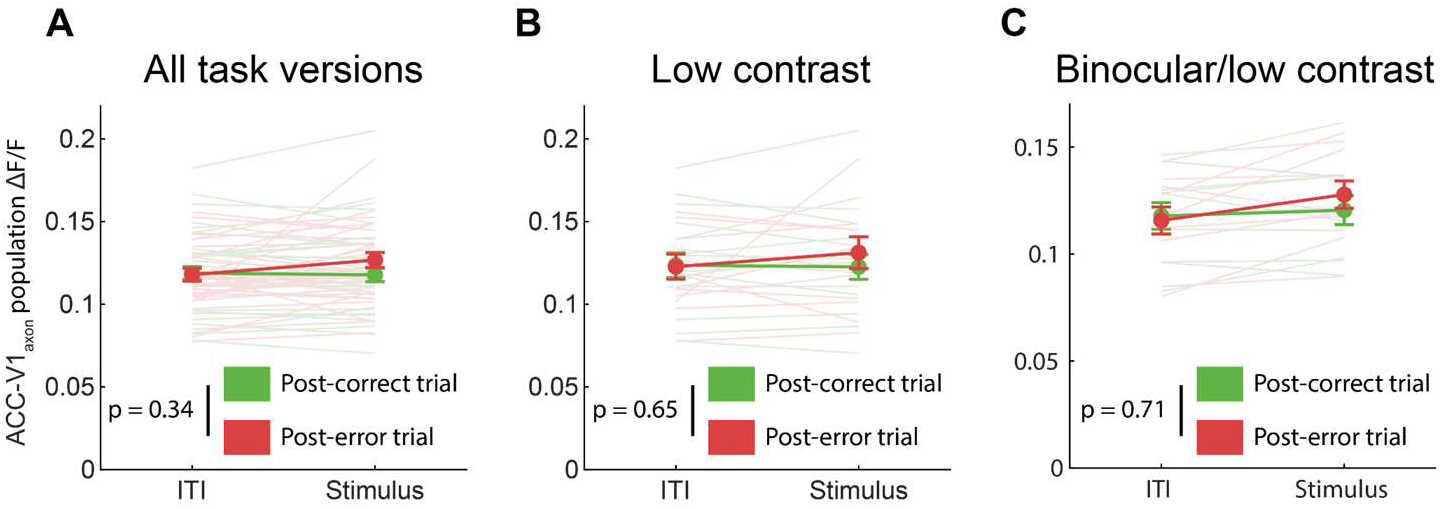
Post-error recruitment of ACC-V1_axons_ is not observed under the conditions of our study. (A) Comparison of mean ΔF/F of boutons in each experiment posterror or post-correct trial, either during the intertrial period (ITI) or during the stimulus period of the trial. All experiments are included in analysis. (B) As in (A) for low contrast experiments. (C) As in (A) but for bilateral low contrast experiments (arguably the most difficult condition).

